# No robust coexistence in a canonical model of plant-soil feedbacks

**DOI:** 10.1101/2021.10.27.466177

**Authors:** Zachary R. Miller, Pablo Lechón-Alonso, Stefano Allesina

## Abstract

Plant-soil feedbacks (PSFs) are considered a key mechanism generating frequency-dependent dynamics in plant communities. Negative feedbacks, in particular, are often invoked to explain coexistence and the maintenance of diversity in species-rich communities. However, the primary modeling framework used to study PSFs considers only two plant species, and we lack clear theoretical expectations for how these complex interactions play out in communities with natural levels of diversity. Here, we demonstrate that this canonical model for PSFs is equivalent to a well-studied model from evolutionary game theory, and use this equivalence to characterize the dynamics with an arbitrary number of plant species. Surprisingly, we find that coexistence of more than two species is virtually impossible, suggesting that alternative theoretical frameworks are needed to describe feedbacks observed in diverse natural communities. Drawing on our analysis, we discuss future directions for PSF models and implications for experimental study of PSF-mediated coexistence in the field.

## 1 Introduction

It has become well understood that reciprocal interactions between plants and the soil biota, collectively termed plant-soil feedbacks (PSFs), play an important role in structuring the composition and dynamics of plant communities. PSFs operate alongside other factors, including abiotic drivers (Bennett & Klironomos 2019) and above-ground trophic interactions (Van der Putten *et al.* 2009), but are thought to be a key mechanism generating negative frequency-dependent feedbacks that promote coexistence and maintain plant diversity *(Kulmatiski et al.* 2008; Van der Putten *et al.* 2013; Bever *et al.* 2015). The existence of PSFs has long been known (Van der Putten *et al.* 1993; Bever 1994), but our understanding of their importance – particularly in relation to patterns of coexistence – has developed rapidly in recent years (Klironomos 2002; Petermann *et al.* 2008; Mangan *et al.* 2010; Crawford *et al.* 2019). Broad interest in PSFs was ignited by the development of simple mathematical models, which illustrated the potential of PSFs to mediate plant coexistence (Bever *et al.* 1997; Bever 2003; Ke & Miki 2015). These models have played a crucial guiding role for a wide range of empirical studies, as well (Kulmatiski *et al.* 2008; Pernilla Brinkman *et al.* 2010; Kulmatiski *et al.* 2011).

The first, and still most widely known and used, model for PSFs was introduced by Bever and colleagues in the 1990s (Bever 1992; Bever *et al.* 1997; Bever 1999). In this framework, often referred to simply as the Bever model, each plant species is assumed to promote the growth of a specific soil component (i.e. associated bacteria, fungi, invertebrates, considered collectively) in the vicinity of individual plants. In turn, the fitness of each plant species is determined by the relative frequency of different soil components. Starting from minimal assumptions, Bever *et al.* (1997) derived a set of differential equations to capture these dynamics. PSFs can be either positive (fitness of a plant species is higher in soil conditioned by conspecifics, compared to heterospecific soil) or negative (a plant species experiences lower relative fitness in its own soil). Bever et *al.* introduced a single quantity to summarize whether community-wide PSFs are positive or negative, and showed that this value characterizes the dynamical behavior of the model. In the original Bever model of two plant species, positive PSFs lead to priority effects, and consequently the exclusion of one species, while negative PSFs result in neutral oscillations. It is thus widely suggested that negative PSFs help sustain coexistence in real-world plant communities (Kulmatiski *et al.* 2008; Van der Putten *et al.* 2013), perhaps with spatial asynchrony playing a role in stabilizing the cyclic dynamics (Bever 2003; Revilla *et al.* 2013).

Subsequent studies have generalized PSF models to include, for example, more realistic functional forms (Umbanhowar & McCann 2005; Eppinga *et al.* 2006), more explicit representations of the soil community (Bever *et al.* 2010), spatial structure (Molofsky *et al.* 2002; Eppinga *et al.* 2006; Suding *et al.* 2013), or additional processes such as direct competitive interactions between plants (Bever 2003). However, the original Bever model remains an important touchstone for the theory of PSFs (Ke & Miki 2015; Ke & Wan 2020; Abbott *et al.* 2021) and informs empirical research through the interaction coefficient, *I_s_,* derived by Bever et *al.,* which is commonly measured and used to draw conclusions about coexistence in experimental studies. Despite the ubiquity of this model, and the fruitful interplay of theory and experiment in the PSF literature, extensions to communities with more than two or three species have appeared only rarely and recently (but see Kulmatiski *et al.* 2011; Eppinga *et al.* 2018; Mack *et al.* 2019). While PSF models motivate hypotheses and conclusions about species-rich natural communities, there is much still unknown about the behavior of these models with natural levels of diversity (Van der Putten *et al.* 2013).

Here, we extend the Bever model to include any number of plant species, and show that the model is equivalent to a special form of the replicator equation studied in evolutionary game theory (Hofbauer & Sigmund 1998). In particular, this model corresponds to the class of bimatrix games, where there are two players (here, plants and soil components) which interact with asymmetric strategies and payoffs. The replicator dynamics of bimatrix games are well-studied, allowing us to characterize many properties of the Bever model with *n* plant species. Surprisingly, using this equivalence, we show that coexistence of more than two species in this model is never robust.

## 2 Results

### 2.1 Generalizing a classic PSF model

Inspired by emerging empirical evidence for the important role of PSFs in plant community dynamics and coexistence (Van der Putten *et al.* 1993; Bever 1994), Bever *et al.* (1997) introduced a simple mathematical model to investigate their behavior. In this model, two plant species, 1 and 2, grow exponentially with growth rates determined by the state of the soil biota in the system. These effects of soil on plants are specified by parameters *α_ij_*, the growth rate of plant species *i* in soil type *j*. There is a soil component corresponding to each plant species, which grows exponentially in the presence of its associated plant at a rate *β_i_*. Bever *et al.* set an important precedent by considering the dynamics of *relative* abundances in such a system; starting from dynamics of the form

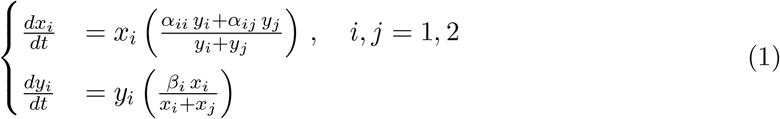

for the *absolute* abundances of plants (*x_i_*) and soil components (*y_i_*), one derives dynamics for the relative abundances (frequencies), *p_i_* = *x_i_* /∑_*j*_ *x_j_* and *q_i_* = *y_i_* /∑_*j*_ *y_j_*. Using the facts *p_i_* = 1 – *p_j_* and *q_i_* = 1 – *q_j_*, these can be written as:

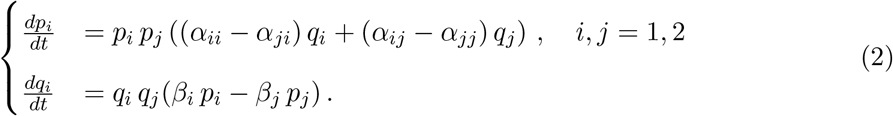

This model may admit a coexistence equilibrium where

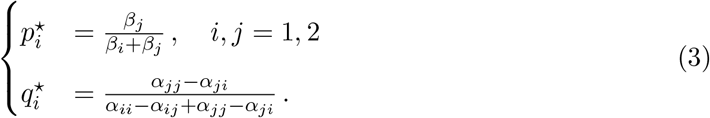

A central finding of the analysis by Bever *et al.* was that the denominator of 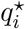, which they termed the “interaction coefficient”, *I_s_* (= *α_ii_* – *α_ij_* + *α_jj_* – *α_ji_*, controls the model dynamics: When *I_s_* > 0, which represents a community with positive feedbacks, the equilibrium in Eq. 3 is unstable, and the two species cannot coexist. On the other hand, when *I_s_* < 0, the equilibrium is neutrally stable, and the dynamics cycle around it, providing a form of nonequilibrium coexistence. In fact, these conclusions also depend on the existence of a feasible equilibrium (i.e. positive equilibrium values), which further requires that *α_ii_* < (*j* for both *i, j* = 1,2, in order for the model to exhibit coexistence (Bever *et al.* 1997; Ke & Miki 2015).

This coexistence is fragile. Plant and soil frequencies oscillate neutrally, similar to the text-book example of Lotka-Volterra predator-prey dynamics. Any stochasticity, external forcing, or time variation in the model parameters can destroy these finely balanced oscillations and cause one species to go extinct (Revilla *et al.* 2013). However, coupled with mechanisms that buffer the system from extinctions, such as migration between desynchronized patches or the presence of a seed bank, the negative feedbacks in this model might produce sustained coexistence (Bever 2003; Revilla *et al.* 2013).

Of course, most natural plant communities feature more than two coexisting species, and it is precisely in the most diverse communities that mechanisms of coexistence hold the greatest interest (Van der Putten *et al.* 2013). While it is not immediately clear how to generalize Eq. 2 to more than two species, Eq. 1 is naturally extended by maintaining the assumption that the overall growth rate for any plant is a weighted average of its growth rate in each soil type:

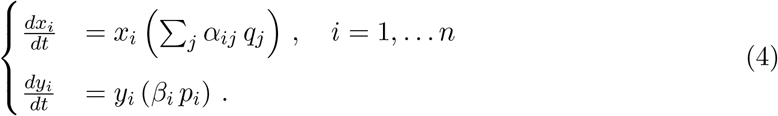

From Eq. 4, one can derive the *n*-species analogue of Eq. 2,

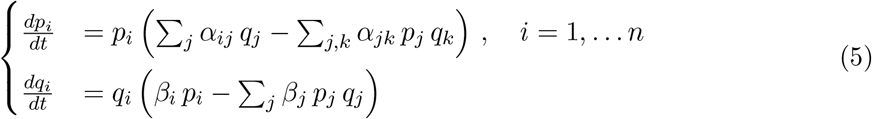

giving the dynamics for species and soil component frequencies (Fig. 1). Eq. 5 is conveniently expressed in matrix form as

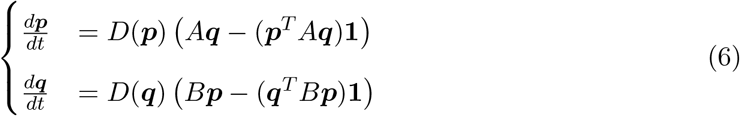

where vectors are indicated in boldface (e.g. ***p*** is the vector of plant species frequencies (*p*_1_, *p*_2_,…*P_n_*)^*T*^ and **1** is a vector of *n* ones) and *D*(***z***) is the diagonal matrix with vector ***z*** on the diagonal. We have introduced the matrices *A* = (*α_ij_*) and *B* = *D*(*β*_1_, *β*_2_,…*β_n_*), specifying soil effects on plants and plant effects on soil, respectively. Because ***p*** and ***q*** are vectors of frequencies, they must sum to one: **1**^*T*^***p*** = **1**^*T*^***q*** = 1. Using these constraints, one can easily show that the Bever model (Eq. 2) is a special case of Eqs. 5 and 6 when *n* = 2.

**Figure 1:**
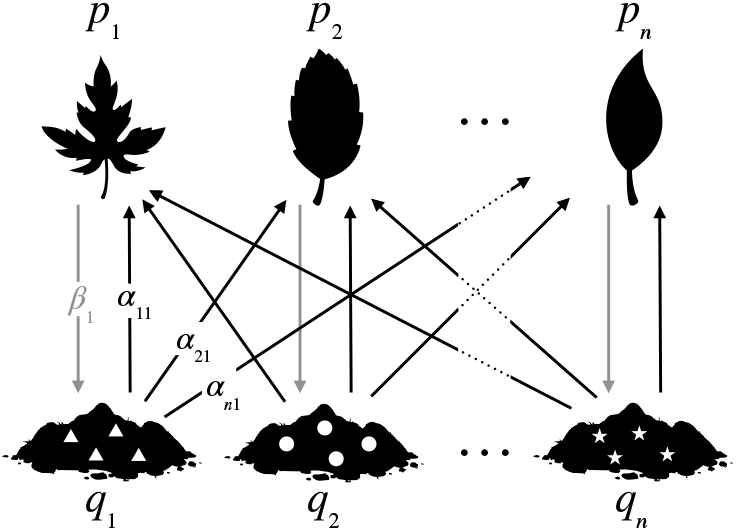
The model described by Eqs. 5-6 is shown here graphically. Plant species (top) promote the growth of their respective soil components (bottom) at a rate *β_i_* (gray arrows). In turn, growth of each plant is governed by the mix of soil components present in the system, with the effect of soil component *j* on species *i* quantified by the parameter *α_ij_* (black arrows). This model is a straightforward extension of the model proposed by Bever *et al.* (1997) to an arbitrary number of species. Note only selected parameter labels are shown for clarity.

This model (Eq. 5) is a direct multispecies generalization of the classic Bever model (Bever *et al.* 1997); it requires precisely the same assumptions as the original, and includes the same biological processes. Other extensions of the Bever model have been introduced, including versions that incorporate plant-plant competition and self-regulation (Bever 2003) or modified soil dynamics (Eppinga *et al.* 2018; Mack *et al.* 2019). Contrasting the predictions of these models can illuminate the connections between particular biological processes (model assumptions) and resulting community dynamics, a point we return to in the Discussion. However, it is also important to recognize that different biological assumptions can give rise to identical dynamics; for example, we show in the Supplemental Methods that the model introduced by Bever in 2003, which includes more realistic plant dynamics and interactions, reduces to our Eq. 5 when plants are competitively equivalent.

### 2.2 Equivalence to bimatrix game dynamics

Systems that take the form of Eq. 6 are well-known and well-studied in evolutionary game theory. Our generalization of the Bever model is a special case of the *replicator equation,* corresponding to the class of *bimatrix games* (Taylor 1979; Hofbauer 1996; Hofbauer & Sigmund 1998; Cressman & Tao 2014). Bimatrix games arise in diverse contexts, such as animal behavior (Taylor 1979; Selten 1988), evolutionary theory (Hofbauer & Sigmund 1998; Cressman & Tao 2014), and economics (Friedman 1991), where they model games with asymmetric players, meaning that each player (here, the plant community and the soil) has a distinct set of strategies (plants species and soil components, respectively) and payoffs (realized growth rates).

Much is known about bimatrix game dynamics, and we can draw on this body of knowledge to characterize the behavior of the Bever model with *n* species. Essential mathematical background and details are presented in the Supplemental Methods; for a detailed introduction to bimatrix games, see Hofbauer & Sigmund (1998).

Under the mild condition that matrix *A* is invertible, Eq. 6 admits a unique coexistence equilibrium given by ***p***\* = *k_p_* B^-1^**1** and ***q***\* = *k_q_ A*^-1^**1**, where *k_p_* = 1/(**1**^*T*^*B*^-1^**1**) and *k_q_* = 1/(**1**^*T*^*A*^-1^**1**) are constants of proportionality that ensure the equilibrium frequencies sum to one for both plants and soil. Because B is a diagonal matrix, and all *β_i_* are assumed positive, the equilibrium plant frequencies, ***p***\*, are always positive, as well. Thus, feasibility of the equilibrium hinges on the soil frequencies, ***q***\*, which are all positive if the elements of *A*^-1^**1** all share the same sign.

As we have seen, when the community consists of two plant species, the coexistence equilibrium, if feasible, can be either unstable or neutrally stable. The same is true for the *n*-species extension (and, more generally, for any bimatrix game dynamics; see Eshel *et al.* 1983; Selten 1988; Hofbauer & Sigmund 1998). This can be established using straightforward local stability analysis, after accounting for the relative abundance constraints, which imply 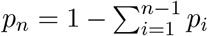 and 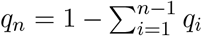. Using these substitutions, Eq. 5 can be written as a system of 2*n* – 2 (rather than 2*n*) equations, and the community matrix for this reduced model has a very simple form (see Supplemental Methods). In particular, the community matrix has all zero diagonal elements, which implies that the eigenvalues of this matrix sum to zero. These eigenvalues govern the stability of the coexistence equilibrium, and this property leaves two qualitatively distinct possibilities: either the eigenvalues have a mix of positive and negative real parts (in which case the equilibrium is unstable), or the eigenvalues all have zero real part (in which case the equilibrium is neutrally stable). Already, we can see that the model never exhibits equilibrium coexistence, regardless of the number of species.

In fact, these conclusions apply to a much broader class of models that are structurally similar to the Bever model. Consider any PSF dynamics that can be written in the form

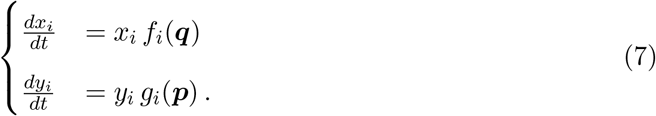

Such a model has a bipartite structure, meaning that the per capita growth rate of each plant species is an arbitrary function, *f_i_*, of the soil frequencies, with no dependence on the plant abundances or frequencies, and vice versa. The corresponding dynamics for plant and soil frequencies may possess one or more coexistence equilibria. At any of these equilibrium points, the community matrix, which governs its stability, necessarily takes the same form as for the Bever model, with all zero diagonal elements. As such, these equilibria must be either unstable or neutrally stable. This is a robust consequence of the strictly bipartite structure of Eq. 7.

Another notable property of bimatrix game dynamics is that the vector field defined by the model equations is divergence-free or incompressible (see Hofbauer & Sigmund 1998, for a proof). The divergence theorem from vector calculus (Arfken 1985) then dictates that Eq. 6 cannot have any attractors – that is, regions of the phase space that “pull in” trajectories – with multiple species. This rules out coexistence in a stable limit cycle or other nonequilibrium attractors (e.g. chaotic attractors). Thus, only the relatively fragile coexistence afforded by neutral oscillations is possible, as in the two-species model.

Based on the local stability properties of the coexistence equilibrium, Bever *et al.* concluded that such neutral cycles arise for two species when an *α*_11_ < *α*_21_ and *α*_22_ < *α*_42_. The equivalence between their model and a bimatrix game with two strategies allows us to give a fuller picture of these cycles. Namely, we can identify a constant of motion for the two-species dynamics:

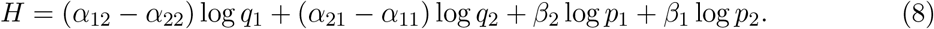

Using the chain rule and time derivatives in Eq. 2, it is easy to show that 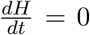 for any plant and soil frequencies (see Supplemental Methods). The level curves of *H* form closed orbits around the equilibrium when the equilibrium is neutrally stable. Thus, *H* implicitly defines the trajectories of the model, and can be used to determine characteristics such as the amplitude of oscillations arising from particular initial frequencies (Volterra 1926).

Because neutral cycles provide the only possible form of coexistence in this model, a key question becomes whether and when neutral cycles with *n* plant species can arise. Do the “negative feedback” conditions identified by Bever *et al.* generalize in richer communities? Indeed, they do; however, for more than two species, these conditions are very severe. The model in Eq. 6 supports oscillations with *n* plant species – for any *n* – if matrices *A* and *B* satisfy a precise relationship (see Supplemental Methods for details). In particular, the model parameters must satisfy the conditions *α_ij_* = *γ_i_* + *δ_j_* for some constants *γ_i_,δ_i_* in *i* = 1,…, *n* (when *i* ≠ *j*), and *α_ii_* = *γ_i_* + *δ_i_* – *cβ_i_* (where *c* is a positive constant that must be the same for all species *i*). In the language of bimatrix games, such systems are called *rescaled zerosum games* (Hofbauer 1996; Hofbauer & Sigmund 1998). It is a long-standing conjecture in evolutionary game theory that these parameterizations are the *only* cases where *n*-species coexistence can occur (Hofbauer 1996, 2011).

Ecologically, these conditions mean there is a fixed effect of each soil type and plant species identity, and the growth rate of plant *i* in soil type *j* is the additive combination of these two, with no interaction effects. The only exception is for plants growing in their own soil type, which must experience a fitness cost (*γ_i_* + *δ_i_* – *α_ii_*) exactly proportional to the rate at which they promote growth of their soil type (*β_i_*). These conditions clearly extend the intuitive notion that each plant must have a disadvantage in its corresponding soil type to allow for coexistence. But the parameters of the model are constrained so strongly that we never expect to observe cycles with more than two plant species in practice. When *n* > 2, a great deal of fine-tuning is necessary to satisfy the rescaled zero-sum game condition; the probability that random parameters will be suitable is infinitesimally small. We confirm this numerically with simulations shown in Fig. 2. Although *n*-species cycles are clearly possible (as in Fig. 3), when parameters are drawn independently at random communities always collapse to one or two species, regardless of the initial richness.

**Figure 2:**
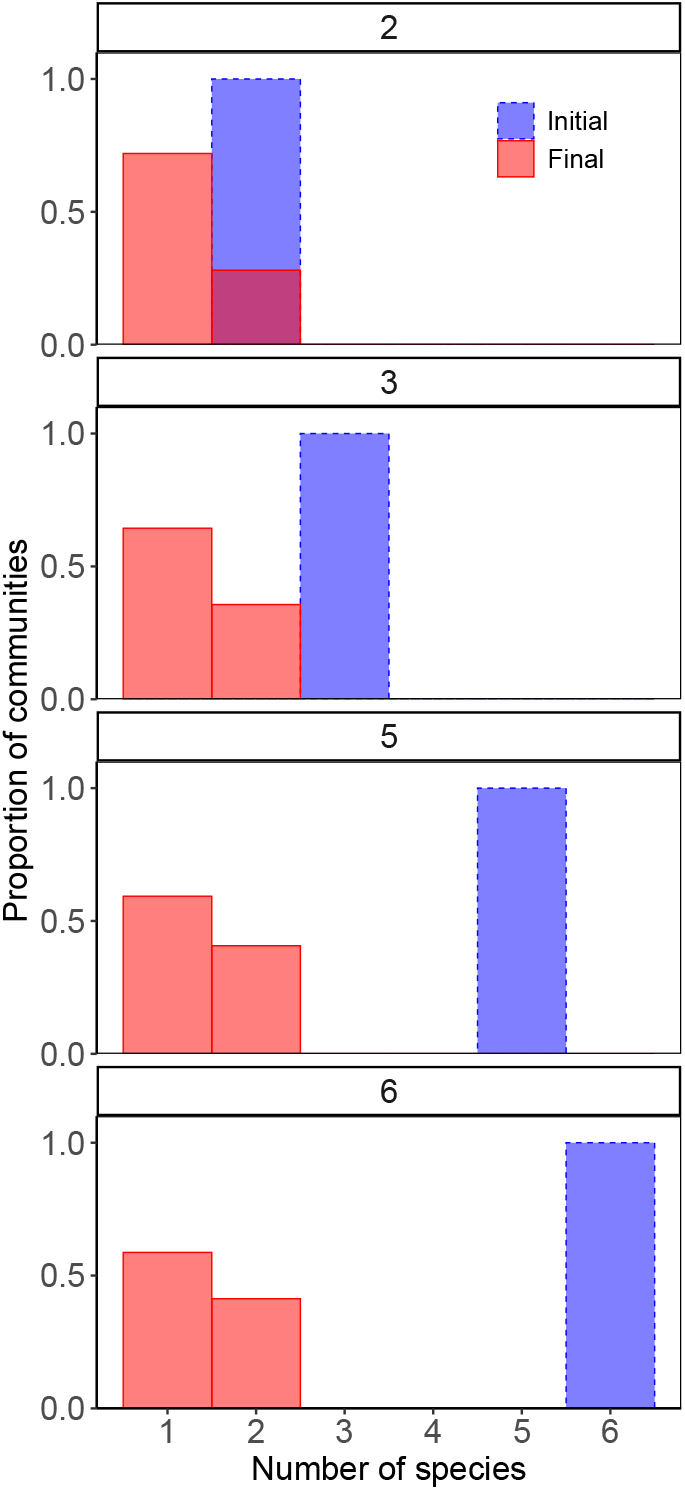
Final community sizes with varying initial richness. We show the distribution of final richness (number of species, in red) for 5000 communities governed by the *n*-species Bever model, initialized with 2, 3, 5, or 6 plant species. Parameters *α_ij_* and *β_i_* were sampled independently from a standard uniform distribution, *U*(0,1). For each random parameterization at each level of initial richness, we integrated the dynamics of Eq. 6 until the system reached a periodic orbit or until only one species remained. In agreement with the conjecture that coexistence of more than two species is vanishingly unlikely, we found that regardless of the number of species initially present, every community collapsed to a subset with one or two surviving species.

**Figure 3:**
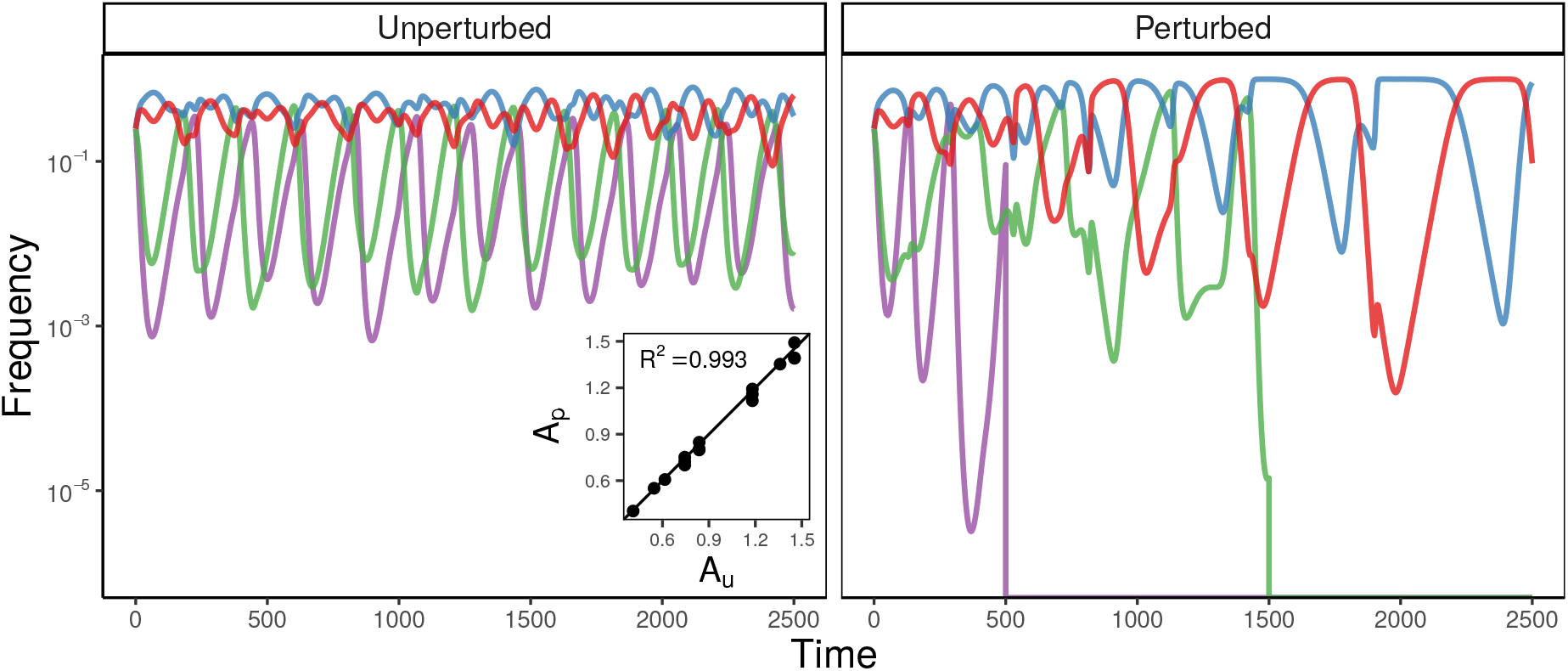
Coexistence of three or more species is not robust. It is possible to obtain neutrally stable oscillations with any number of plant species if the model parameters constitute a rescaled zero-sum game (see text for conditions). Here, for example, we show sustained oscillations with 4 plant species (soil frequencies not shown) using fine-tuned parameters, *A_u_* (left). However, if we randomly perturb *A_u_* by a small amount to obtain new parameters, *A_p_*, the dynamics quickly collapse to a two-species subset (right). Any slight perturbation is enough to disrupt coexistence; for this example, the parameters *A_u_* and *A_p_* are highly correlated (inset) and differ in value by less than 3% on average.

Parameter combinations permitting many-species oscillations are not only rare, they are also extremely sensitive to small changes to the parameter values. The rescaled zero-sum condition imposes many exact equality constraints on the matrix *A* (e.g. *α_ij_* – *α_ik_* = *α_lj_* – *α_lk_* for all *i,j,k,* and *l*). Even if biological mechanisms exist to generate the requisite qualitative patterns, inevitable quantitative variation in real-world communities will disrupt coexistence (Fig. 3). Coexistence of *n* > 2 plant species – even in the weak sense of neutral cycles – is not robust to small changes in the model parameters.

Interestingly, the two-species model is not subject to the same fragility. It can be shown (see Supplemental Methods) that all 2 × 2 bimatrix games take the same general form as a rescaled zero-sum game, although the constant *c* may be positive or negative, depending on the parameters. When *I_s_*, the interaction coefficient identified by Bever *et al.,* is negative, c is positive, ensuring (neutral) stability. This condition amounts to an inequality constraint, rather than an equality constraint, and so it *is* generally robust to small variations in model parameters. As we can now see, the case *n* = 2 is unique in this regard.

## 3 Discussion

The Bever model has played a central role in motivating PSF research, and continues to guide both theory and experiment in this fast-growing field (Bever *et al.* 2015; Kandlikar *et al.* 2019; Ke & Wan 2020; Abbott *et al.* 2021). Here, we extend the Bever model to include any number of plant species, and highlight its equivalence to bimatrix game dynamics. Taking advantage of the well-developed theory for these dynamics, we are able to characterize the behavior of this generalized Bever model in detail.

Our central finding is that there can be no robust coexistence of plant species in this model. Regardless of the number of species, *n*, the model never exhibits equilibrium coexistence or other attractors. Coexistence can be attained through neutral oscillations, but these dynamics lack any restoring force, meaning diversity would quickly be eroded by stochasticity or exogenous forcing. In this respect, the generalized model behaves similarly to the classic two-species system. However, unlike the two-species model, oscillations with *n* > 2 species can only occur under very restricted parameter combinations. These parameterizations are vanishingly unlikely to arise by chance and highly sensitive to small deviations. Thus, coexistence of more than two species is neither dynamically nor structurally stable.

This result may seem surprising, because a significant body of experimental evidence indicates that PSFs can and do play an important role in mediating the coexistence of more than two species in natural communities (Kulmatiski *et al.* 2008; Petermann *et al.* 2008; Mangan *et al.* 2010; Bever *et al.* 2015). Apparently, the picture suggested by the two-species Bever model generalizes in nature, but not in the model framework itself. We note that this framework was introduced as an intentional simplification to illustrate the potential role of PSFs in mediating coexistence, not to accurately model the biological details of PSFs. Indeed, the model has been wildly successfully in spurring research into PSFs. Alongside extensive empirical study of these processes, other modeling approaches have emerged, accounting for more biological realism (e.g., Umbanhowar & McCann 2005; Eppstein & Molofsky 2007; Bever *et al.* 2010), or with the demonstrated capacity to produce multispecies coexistence (e.g., Bonanomi *et al.* 2005; Eppinga *et al.* 2018; Miller & Allesina 2021). Some of these are minor modifications of the Bever model framework; others build on distinct foundations (Ke & Miki 2015; Ke & Wan 2020). Our results suggest that these various avenues are worth pursuing further.

Our findings also help clarify important aspects of coexistence across modeling approaches. For example, Eppinga *et al.* (2018) and Mack *et al.* (2019) recently introduced a multispecies PSF model which can exhibit stable coexistence. Their model is inspired by the Bever model framework, but departs from it in two ways: by introducing more realistic soil dynamics, including a carrying capacity for soil communities, and by applying a separation of timescales, under the assumption that soil dynamics are very rapid compared to plant dynamics. Our analysis indicates that the second feature is unable to account for stabilization. Regardless of the relative rates of plant and soil dynamics, the coexistence equilibrium of the generalized Bever model is never attractive. This is a fundamental feature of the model structure, not a result of particular parameter choices. When oscillations do exist in the Bever model, they are always neutral, meaning that their amplitude is fixed by the initial conditions of the system, and cannot diminish through the dynamics. These observations make clear that the crucial factor driving coexistence in the model of Eppinga, Mack, and colleagues is self-regulation within soil communities, not rapid soil dynamics.

Indeed, our analysis suggests that the internal dynamics of plant or soil communities must interact with PSFs to maintain diversity in natural systems. We have shown that PSF models that are structurally similar to the Bever model – in which plant dynamics depend only on soil frequencies, and soil dynamics depend only on plant frequencies – are incapable of exhibiting stable coexistence of any number of species. Multispecies coexistence becomes possible when plants (Bever 2003; Revilla *et al.* 2013), soils (Eppinga *et al.* 2018; Mack *et al.* 2019), or both experience an independent source of self-regulation, which might arise from resource competition, physical limits to density, or some other mechanism. PSFs are likely to matter most for the maintenance of diversity when they interact with these internal plant or soil dynamics in non-trivial ways. In the case of combined plant competition-feedback models (Bever 2003), for example, we have already seen that no robust coexistence is possible when plants are competitively equivalent and experience “mean-field” interactions. On the other hand, when plant competition is dominated by strong intraspecific interactions, all plant species would coexist even in the absence of PSFs. Thus, PSFs can only contribute to the maintenance of diversity in such models by modifying competitive outcomes (Bever 2003; Revilla *et al.* 2013; Kandlikar *et al.* 2019) – that is, by interacting with the structure of the plant-plant competitive network.

These conclusions have practical implications for the study of PSFs in real-world communities. The predictions of the Bever model are commonly used to guide the design and analysis of PSF experiments, especially in drawing conclusions about coexistence. Our analysis cautions that direct application of this model in multispecies communities might lead to incorrect inference. For example, attempts to parameterize the Bever model for three species using empirical data have produced predictions of non-coexistence in plant communities that coexist experimentally (Kulmatiski *et al.* 2011). In many other studies, the interaction coefficient, *I_s_,* is calculated for species pairs and used to assess whole-community coexistence (Kulmatiski *et al.* 2008; Fitzsimons & Miller 2010; Pendergast IV *et al.* 2013; Suding *et al.* 2013; Kuebbing *et al.* 2015; Smith & Reynolds 2015; Bauer *et al.* 2017; Pizano *et al.* 2019; Crawford *et al.* 2019). However, we have seen that whole-community coexistence is virtually impossible within the generalized model, and there is no guarantee that the pairwise coexistence conditions for this model will agree with *n*-species coexistence conditions in other frameworks. For example, *I_s_* < 0 for all species pairs is neither necessary nor sufficient to produce coexistence in a metapopulation-based model for PSFs (Miller & Allesina 2021).

Theory suggests that when PSFs do play a role in maintaining robust coexistence, interactions between plants and soil will necessarily be only part of the picture. On this point, we echo calls that have emerged in the empirical literature for more closely integrated study of PSFs and other processes, such as plant competition (Casper & Castelli 2007; Lekberg *et al.* 2018) and more detailed soil biology (Bever *et al.* 2010; Hodge & Fitter 2013). Our results strongly suggest that pairwise PSF measurements are insufficient to characterize plant coexistence and require contextualization alongside these other ecosystem processes.

Fundamentally, our analysis demonstrates that PSFs as envisioned in the classic Bever model cannot produce robust *n*-species coexistence in isolation. Our results also indicate basic structural features that are necessary for PSF models to support multispecies coexistence. Significantly, we find not only the absence of stabilization in the Bever model, but generic *instability.* This suggests that, in diverse communities, other processes must exert a sufficiently strong influence on the community dynamics to overcome the baseline instability. We illustrate this idea in the Supplemental Methods, where we examine the effect of adding negative frequency-dependence in the classic Bever model. Any amount of frequency-dependence stabilizes neutral oscillations, but when these effects are weak, they cannot turn unstable equilibria into stable ones. The result is that the augmented model can only support multispecies coexistence when the rescaled zero-sum game condition is met, and, as we have shown, this condition is never robust to small parameter variations.

In this example, we consider negative frequency-dependence, rather than density-dependence (as in the well-known Bever (2003) model), because it is difficult to compare the strengths of processes that mix units of frequency and density. This difficulty hints at a central limitation of classic PSF models, which are derived as projections (onto the space of relative abundances, or frequencies) of dynamics for plant and soil *abundances* (Kulmatiski *et al.* 2011; Revilla *et al.* 2013; Eppinga *et al.* 2018; Ke & Wan 2020). The projected dynamics can mask unbiological outcomes in the original model (e.g. relative abundances oscillate around equilibrium while absolute abundances shrink to zero or explode to infinity). Indeed, the absolute abundance model (Eq. 4) used to derive our *n*-species frequency dynamics (Eqs. 5-6) does not generally possess any fixed point, which is a basic requirement for species coexistence (Hutson 1990; Hutson & Schmitt 1992). The same is true for the model introduced by Eppinga *et al.* (2018) and Mack *et al.* (2019), even though this model exhibits stable dynamics for plant and soil frequencies. It is usually seen as desirable to study PSFs in the space of species frequencies, both because this facilitates connections to data, and because frequencies are considered a more appropriate metric for analyzing processes that stabilize coexistence (Adler *et al.* (2007); Eppinga *et al.* (2018), but see Kandlikar *et al.* (2019); Ke & Wan (2020)). But models that introduce frequencies through a natural constraint, such as competition for finite space, will likely produce more realistic and more straightforwardly interpretable dynamics.

From a broader theoretical perspective, the qualitative change in model behavior that we observe as the number of species increases from two to three or more is a striking phenomenon, but not an unprecedented one. Ecologists have repeatedly found that intuitions from two-species models can generalize (or fail to generalize) to more diverse communities in surprising ways (Strobeck 1973; Smale 1976; Barábas *et al.* 2016). Our analysis provides another illustration of the fact that “more is different” (Anderson 1972) in ecology, and highlights the importance of developing theory for species-rich communities.

## Acknowledgments

We thank Joy Bergelson for helpful comments and two anonymous reviewers for feedback that improved and broadened the text.

## Data accessibility

Code for reproducing the numerical simulations is available at https://github.com/pablolich/plant_soil_feedback.

## Notes

https://github.com/pablolich/plant_soil_feedback

### Competing Interest Statement

The authors have declared no competing interest.

