## Supplemental Methods for "No robust coexistence in a canonical model of plant-soil feedbacks"

#### 1 Model derivation

As described in the Main Text, we begin with the system

$$\begin{cases} \frac{dx_i}{dt} & = x_i \left( \sum_j \alpha_{ij} \, q_j \right) , \quad i = 1, \dots n \\ \frac{dy_i}{dt} & = y_i \left( \beta_i \, p_i \right) \end{cases} \tag{S1}$$

governing the time-evolution of plant abundances $x_i$ and soil components $y_i$, where $p_i = x_i / \sum_j x_j$, $q_i = y_i / \sum_j y_j$, and Greek letters denote nonnegative parameters. These equations capture the assumptions outlined by Bever *et al.* (1997) for two species and extend them straightforwardly to any $n$ species. Following the approach of Bever *et al.* for two species (and consistent with other generalizations of this model, e.g., Kulmatiski *et al.* 2008; Eppinga *et al.* 2018), we derive dynamics for frequencies by applying the chain rule:

$$
\begin{aligned}
\frac{dp_i}{dt} &= \frac{d}{dt}\frac{x_i}{\sum x_j} \\
&= \frac{1}{\sum_j x_j}\frac{dx_i}{dt} - \frac{x_i}{(\sum_j x_j)^2}\sum_j \frac{dx_j}{dt} \\
&= \frac{x_i}{\sum_j x_j}\left(\sum_j \alpha_{ij}\, q_j\right) - \frac{x_i}{\sum_j x_j}\left(\sum_j \frac{x_j}{\sum_k x_k}\sum_l \alpha_{jl}\, q_l\right) \\
&= p_i\left(\sum_j \alpha_{ij}\, q_j - \sum_{j,k} \alpha_{jk} p_j q_k\right) .
\end{aligned}
\tag{S2}
$$

This last expression is identical to the first line of Eq. 5 in the Main Text. The dynamics for $q_i$ can be derived in exactly the same way (using the definitions $\beta_{ii} = \beta_i$ and $\beta_{ij} = 0$). The two terms of each per capita growth rate in Eq. 5 have natural interpretations in the language and notation of linear algebra: $\sum_j \alpha_{ij}\, q_j$ is the $i$th component of the matrix-vector product $A\boldsymbol{q}$ and $\sum_{j,k} \alpha_{jk} p_j q_k$ is the bilinear form $\boldsymbol{p}^T A\boldsymbol{q}$. Here, $A$ (and $B$) is an $n \times n$ matrix and $\boldsymbol{p}$ and $\boldsymbol{q}$ are vectors of length $n$, as described in the Main Text. We can re-write Eq. 5 as

$$
\begin{aligned}
\frac{dp_i}{dt} &= p_i\left((A\boldsymbol{q})_i - \boldsymbol{p}^T A\boldsymbol{q}\right) \\
\frac{dq_i}{dt} &= q_i\left((B\boldsymbol{p})_i - \boldsymbol{q}^T B\boldsymbol{p}\right)
\end{aligned}
\tag{S3}
$$

or even more compactly as

$$
\begin{cases}
\frac{d\boldsymbol{p}}{dt} &= D(\boldsymbol{p})\left(A\boldsymbol{q} - (\boldsymbol{p}^T A\boldsymbol{q})\mathbf{1}\right) \\
\frac{d\boldsymbol{q}}{dt} &= D(\boldsymbol{q})\left(B\boldsymbol{p} - (\boldsymbol{q}^T B\boldsymbol{p})\mathbf{1}\right)
\end{cases}
\tag{S4}
$$

which is Eq. 6 in the Main Text.

An alternative derivation of these dynamics (Eqs. 5 and 6) takes the model introduced by Bever (2003) as a starting point. Using our notation, this model can be written as

$$\begin{cases} \frac{dx_i}{dt} &= x_i \left( r_i + \sum_j \alpha_{ij} \, q_j - \sum_j c_{ij} x_j \right) \ , \quad i = 1, \ldots n \\ \frac{dy_i}{dt} &= y_i \left( \beta_i \, p_i \right) \end{cases} \tag{S5}$$

where all variables have the same meaning as before. In this model, plants experience competitive Lotka-Volterra dynamics alongside frequency-dependent soil effects. The parameters $r_i$ are intrinsic growth rates for plants, and the $c_{ij}$ quantify the competitive effect of plant $j$ on plant $i$, as in the usual Lotka-Volterra model. We note that in this context, the soil effects on plants, $\alpha_{ij}$ may be positive or negative, as they modify the baseline plant growth rates, set by $r_i$. The dynamics of soil communities are exactly as before.

One can write the dynamics for plant frequencies under this model as:

$$\frac{dp_i}{dt} = p_i \left( r_i + \sum_j \alpha_{ij} \, q_j - \sum_j c_{ij} x_j - \sum_j p_j \left[ r_j + \sum_k \alpha_{jk} \, q_k - \sum_k c_{jk} x_k \right] \right) \ , \quad i = 1, \ldots n \tag{S6}$$

following a calculation similar to Eq. S2. As other researchers have noted (Bever 2003; Eppinga *et al.* 2018), if $r_i = r$ and $c_{ij} = c$ for all $i$ and $j$, indicating a situation where all plants are demographically and competitively equal, then Eq. S6 reduces to

$$\frac{dp_i}{dt} = p_i \left( \sum_j \alpha_{ij} \, q_j - \sum_j p_j \sum_k \alpha_{jk} \, q_k \right) \ , \quad i = 1, \ldots n \tag{S7}$$

which is identical to the dynamics for plant frequencies shown in Eq. 5 of the Main Text. Thus, under the simplifying assumption of "mean-field" plant interactions, the two models yield equivalent dynamics for plant and soil frequencies. We will show at the end of this section that the potential difference in signs (i.e. $\alpha_{ij}$ must be nonnegative in the first model formulation, but may take any sign here) has no effect on the dynamics.

The system described by Eq. S4, however obtained, is identical to standard bimatrix replicator dynamics (Hofbauer 1996; Hofbauer & Sigmund 1998). Bimatrix games have two strategy sets (here, the $p_i$ and $q_i$), and interactions take place only between strategies from opposite sets. The growth rate terms we considered above now have interpretations as payoffs or fitnesses: $\sum_j \alpha_{ij}\, q_j = (A\boldsymbol{q})_i$ is the payoff for strategy $i$ (an average of payoffs playing against each strategy of the other "player", weighted by the frequency of each strategy, $q_j$) and $\sum_{j,k} \alpha_{jk} p_j q_k = \boldsymbol{p}^T A \boldsymbol{q}$ is the average payoff across the population of strategies. A general bimatrix game may have any $B$; our model assumptions lead to the special case where $B$ is diagonal. We note that one could easily and plausibly consider an extension of the Bever model where each plant species has some effect on (up to) all $n$ of the soil components. Then, our PSF model would be map exactly onto the full space of bimatrix game dynamics (rather than just a subset). However, all of the results we consider hold for arbitrary bimatrix games, meaning the same conclusions about the dynamics of Eqs. 5-6 would apply to this extended model, as well.

We note two useful properties of Eqs. 5-6, as they will be important for the analysis that follows. First, we have the constraint $\sum_i p_i = \sum_i q_i = 1$ at every point in time. Second, the dynamics are completely unchanged by adding a constant to any *column* of the parameter matrices $A$ or $B$. The first fact is a direct consequence of our definition for $p_i$ and $q_i$; the second can easily be shown. Suppose we have added a constant $w$ to each element in the $l$th column of $A$. Then

$$\begin{aligned}
\frac{dp_i}{dt} &= p_i \left( \sum_j \alpha_{ij}\, q_j + w\, q_l - \sum_{j,k} \alpha_{jk}\, p_j\, q_k - \sum_j w\, p_j\, q_l \right) \\
&= p_i \left( \sum_j \alpha_{ij}\, q_j + w\, q_l - \sum_{j,k} \alpha_{jk}\, p_j\, q_k - w\, q_l \right) \\
&= p_i \left( \sum_j \alpha_{ij}\, q_j - \sum_{j,k} \alpha_{jk}\, p_j\, q_k \right)
\end{aligned} \tag{S8}$$

which is precisely the differential equation we obtained prior to adding $w$. Clearly the trajectories of both systems (with and without the column shift) must be identical. The same considerations apply for the matrix $B$. Intuitively, this property reflects the fact that we are always subtracting the average payoff, and so any change to the payoffs that benefits (or harms) each species equally is "invisible" to the dynamics.

In the remaining sections, we outline the main behaviors of Eqs. 5-6, especially with regard to coexistence. We closely follow the treatment by Hofbauer & Sigmund (1998), and urge interested readers to consult this excellent introduction (see especially chapters 10 and 11). Here, we reproduce or sketch the essential details needed to justify the results in the Main Text.

#### 2 Coexistence equilibrium

Written in matrix form, it is easy to see that the model admits a unique fixed point where all species are present at non-zero frequency. This fixed point, $(\boldsymbol{p}^\star, \boldsymbol{q}^\star)$, must take the form $(k_p B^{-1}\mathbf{1}, k_q A^{-1}\mathbf{1})$ for some undetermined constants $k_p$ and $k_q$. Substituting this ansatz into the growth rates in Eq. 6 and equating them to zero, we have

$$\begin{aligned} A\boldsymbol{q}^\star - ((\boldsymbol{p}^\star)^T A\boldsymbol{q}^\star)\mathbf{1} &= k_q A A^{-1}\mathbf{1} - (k_p k_q \mathbf{1}^T (B^{-1})^T A A^{-1}\mathbf{1})\mathbf{1} = k_q(1 - k_p \mathbf{1}^T (B^{-1})^T \mathbf{1})\mathbf{1} = 0 \\ B\boldsymbol{p}^\star - ((\boldsymbol{q}^\star)^T B\boldsymbol{p}^\star)\mathbf{1} &= k_p B B^{-1}\mathbf{1} - (k_p k_q \mathbf{1}^T (A^{-1})^T B B^{-1}\mathbf{1})\mathbf{1} = k_p(1 - k_q \mathbf{1}^T (A^{-1})^T \mathbf{1})\mathbf{1} = 0 \end{aligned} \tag{S9}$$

From the final two equations, it is clear that $k_p = \frac{1}{\mathbf{1}^T (B^{-1})^T \mathbf{1}} = \frac{1}{\mathbf{1}^T B^{-1}\mathbf{1}}$ and $k_q = \frac{1}{\mathbf{1}^T (A^{-1})^T \mathbf{1}} = \frac{1}{\mathbf{1}^T A^{-1}\mathbf{1}}$.

These rescaling factors make intuitive sense, as they ensure that $\sum_i p_i^\star = \sum_i q_i^\star = 1$, consistent with their definition as frequencies.

Describing these equilibrium frequencies in terms of the parameters is a difficult problem that has received significant attention elsewhere (Eppinga *et al.* 2018; Mack *et al.* 2019; Saavedra *et al.* 2017; Serván *et al.* 2018; Pettersson *et al.* 2020; Saavedra & AlAdwani 2021). In particular, one is usually interested in identifying whether all of the frequencies are non-negative (such a fixed point is said to be feasible). The existence of a feasible fixed point is a requirement for the model to exhibit permanence, meaning that no species go extinct or grow to infinity. Throughout our analysis, we assume the existence of a feasible fixed point; considering the question of feasibility simultaneously would only make coexistence less likely in each case. We present some additional details regarding feasibility in the section *Equilibrium* *feasibility*, below.

#### 3 Local stability analysis

Perturbations around the coexistence equilibrium are constrained to respect the conditions $\sum_i p_i = \sum_i q_i = 1$. For this reason, it is convenient to remove these constraints before performing a local stability analysis. As in the two species case (Bever *et al.* 1997), this can be done by eliminating the $n$th species and soil component, which leaves us with a $2n-2$ dimensional system with no special constraints.

We use $p_n = 1 - \sum_{i=1}^{n-1} p_i \equiv f(\boldsymbol{p})$ and $q_n = 1 - \sum_{i=1}^{n-1} q_i \equiv g(\boldsymbol{q})$ and write these frequencies as functions of the others. The reduced dynamics are given by

$$\begin{cases} \frac{dp_i}{dt} & = p_i \left( \sum_j^{n-1} \alpha_{ij}\, q_j + \alpha_{in} g(\boldsymbol{q}) - \sum_{j,k}^{n-1} \alpha_{jk}\, p_j\, q_k - f(\boldsymbol{p}) \sum_j^{n-1} \alpha_{nj} q_j - g(\boldsymbol{q}) \sum_j^{n-1} \alpha_{jn} p_j - \alpha_{nn} f(\boldsymbol{p}) g(\boldsymbol{q}) \right) \\ \frac{dq_i}{dt} & = q_i \left( \beta_i\, p_i - \sum_j^{n-1} \beta_j\, p_j\, q_j - \beta_n f(\boldsymbol{p}) g(\boldsymbol{q}) \right) , \quad i = 1, \ldots n-1 \end{cases} \tag{S10}$$

Although these equations appear more complex, it is now straightforward to analyze the local stability of the coexistence equilibrium.

The elements of the community matrix (the Jacobian evaluated at the coexistence equilibrium) are easily computed from Eq. S10. First we consider the plant dynamics differentiated with respect to the plant frequencies. In these calculations, all frequencies are evaluated at their equilibrium values.

$$\begin{aligned} \frac{\partial}{\partial p_j} \frac{dp_i}{dt} &= p_i \left( - \sum_k^{n-1} \alpha_{jk}\, q_k + \sum_k^{n-1} \alpha_{nk} q_k - \alpha_{jn} g(\boldsymbol{q}) + \alpha_{nn} g(\boldsymbol{q}) \right) \\ &= p_i \left( - \sum_k^{n} \alpha_{jk}\, q_k + \sum_k^{n} \alpha_{nk} q_k \right) \\ &= 0 \end{aligned} \tag{S11}$$

Here, we have used the fact that $A\boldsymbol{q}^\star \propto \mathbf{1}$. Notice that, because the factors in parentheses in Eq. S10 are zero at equilibrium, these community matrix calculations are valid even for $i = j$.

The other elements are computed similarly:

$$\frac{\partial}{\partial q_j}\frac{dq_i}{dt} = q_i\left(-\beta_i q_i + \beta_n f(\boldsymbol{p})\right) \\ = 0 \tag{S12}$$

$$\frac{\partial}{\partial q_j}\frac{dp_i}{dt} = p_i\left(\alpha_{ij} - \alpha_{in} - \sum_{k}^{n-1}\alpha_{kj}\,p_k - \alpha_{nj}\,f(\boldsymbol{p}) + \sum_{k}^{n-1}\alpha_{kn}p_k + \alpha_{nn}f(\boldsymbol{p})\right) \\ = p_i(\alpha_{ij} - \alpha_{in}) \tag{S13}$$

$$\frac{\partial}{\partial p_j}\frac{dq_i}{dt} = \begin{cases} q_i\,\beta_i\,, & i = j \\ 0\,, & i \neq j \end{cases} \tag{S14}$$

From these calculations, it is apparent that the trace of the community matrix, given by $\sum_i^{n-1}\frac{\partial}{\partial p_i}\frac{dp_i}{dt} + \sum_j^{n-1}\frac{\partial}{\partial q_j}\frac{dq_j}{dt}$ , is zero. The trace of a square matrix is equal to the sum of its eigenvalues (Horn & Johnson 2012), so the eigenvalues of the community matrix must include either (i) a mix of positive and negative real parts or (ii) only purely imaginary values. In the first case, the coexistence equilibrium is locally unstable, because at least one eigenvalue has positive real part. In the second case, the coexistence equilibrium is neutrally or marginally stable. These two possibilities exclude asymptotically stable equilibria. In this respect, the behavior of the two-species model is the generic behavior of the generalized $n$-species model.

### 4 Zero divergence implies no attractors

We can extend this picture beyond a local neighborhood of the coexistence equilibrium by considering the divergence of the vector field associated with Eqs. 5-6. The divergence, defined by $\sum_i \frac{\partial}{\partial p_i}\frac{dp_i}{dt} + \sum_i \frac{\partial}{\partial q_i}\frac{dq_i}{dt}$, measures the outgoing flux around a given point. It can be shown (see Eshel *et al.* 1983; Hofbauer & Sigmund 1998) that up to a change in velocity (i.e., rescaling time by a positive factor), the vector field corresponding to any bimatrix game dynamics has zero divergence everywhere in the interior of the positive orthant (i.e., where $p_i, q_i > 0$ for all

$i$).

The divergence theorem (Arfken 1985) equates the integral of the divergence of a vector field over some $n$-dimensional region to the net flux over the boundary of the region. For a vector field with zero divergence, this implies that every closed surface has zero net flux. As a consequence, such *divergence-free* vector fields cannot have attractors, or subsets of the phase space toward which trajectories of the corresponding dynamical system tend to evolve. If an attractor existed, one could define a surface enclosing it sufficiently tightly, and the net flux over this surface would be negative (as trajectories enter, but do not exit, this region). But this would present a contradiction, and so we conclude that there can be no attractors, such as limit cycles, for the dynamics.

For our model, these facts mean that attractors can only exist on the boundary of the phase space. Because each boundary face for the $n$-dimensional system is another bimatrix replicator system on $n-2$ dimensions, the same logic applies, and the only possible attractors are points where a single species (and corresponding soil component) is present (Hofbauer & Sigmund 1998). States with multiple species present are never attractive. This leaves neutrally-stable oscillations as the only potential form of species coexistence.

#### 5 Rescaled zero-sum games are neutrally stable

In the context of bimatrix games, a zero-sum game is one where $A = -B^T$. A rescaled zero-sum game is one where there exist constants $\gamma_i, \delta_j$ and $c > 0$ such that $a_{ij} + \delta_j = -cb_{ji} + \gamma_i$ for all $i$ and $j$ (here, we understand $A = (a_{ij}), B = (b_{ij})$) (Hofbauer & Sigmund 1998). Any rescaled zero-sum game can be turned into a zero-sum game by adding constants (in particular, $-\delta_j$ and $-\gamma_j$) to each column of $A$ and $B$, and then multiplying $B$ by a positive constant $1/c$. As such, the dynamics of a rescaled zero-sum game and its corresponding zero-sum game are the same up to a rescaling of time.

If a rescaled zero-sum game has a feasible coexistence equilibrium, this equilibrium is neutrally stable. We can see this by considering the associated community matrix. First, we assume without loss of generality that $A = -cB^T$ (otherwise, we shift columns to obtain this form, without altering the dynamics in the process) Now we add the column-constant matrix $\frac{1}{c}\boldsymbol{b}_n\mathbf{1}^T$ to $A$ and $c\boldsymbol{a}_n\mathbf{1}^T$ to $B$, where $\boldsymbol{a}_n$ ($\boldsymbol{b}_n$) denotes the $n$th column of $A$ ($B$). Again, the dynamics, including both equilibrium values and stability properties, are unchanged by this operation. From Eqs. S11-S14, we see that the community matrix, $J$, of the resulting system is given by

$$\begin{pmatrix} 0 & D(\boldsymbol{p}^\star)(\bar{A} + \frac{1}{c}\boldsymbol{b}_n \mathbf{1}^T - \mathbf{1}\boldsymbol{a}_n^T) \\ D(\boldsymbol{q}^\star)(\bar{B} + c\boldsymbol{a}_n \mathbf{1}^T - \mathbf{1}\boldsymbol{b}_n^T) & 0 \end{pmatrix} \tag{S15}$$

where $\bar{A}$ ($\bar{B}$) denotes the $(n-1) \times (n-1)$ submatrix of $A$ ($B$) obtained by dropping the $n$th row and column. Finally, we consider the similarity transform $P^{-1}JP$, defined by the change of basis matrix

$$P = \begin{pmatrix} \sqrt{c}D(\boldsymbol{p}^\star)^{1/2} & 0 \\ 0 & D(\boldsymbol{q}^\star)^{1/2} \end{pmatrix} . \tag{S16}$$

The resulting matrix, $J'$, which shares the same eigenvalues as $J$ (Horn & Johnson 2012), is given by

$$\begin{pmatrix} 0 & \sqrt{c}D(\boldsymbol{p}^\star)^{1/2}(\bar{A} + \frac{1}{c}\boldsymbol{b}_n \mathbf{1}^T - \mathbf{1}\boldsymbol{a}_n^T)D(\boldsymbol{q}^\star)^{1/2} \\ \sqrt{c}D(\boldsymbol{q}^\star)^{1/2}(-\bar{A}^T + \boldsymbol{a}_n \mathbf{1}^T - \frac{1}{c}\mathbf{1}\boldsymbol{b}_n^T)D(\boldsymbol{p}^\star)^{1/2} & 0 \end{pmatrix} \tag{S17}$$

which is a skew-symmetric matrix. Every eigenvalue of a skew-symmetric matrix must have zero real part (Horn & Johnson 2012). Thus, the eigenvalues of $J$, the community matrix, have zero real part, and the coexistence equilibrium of our original system is neutrally stable.

Here, we have outlined a proof that applies to all rescaled zero-sum games. When $B$ is a diagonal matrix, as in our model of PSFs, the condition for $A$ and $B$ to constitute a rescaled zero-sum game reduces to the condition given in the Main Text.

Rescaled zero-sum games are the only bimatrix games known to produce neutrally stable oscillations. It is a long-standing conjecture that no other bimatrix games have this property (Hofbauer 1996; Hofbauer & Sigmund 1998; Hofbauer 2011).

### 6 Two-species bimatrix games

For $n > 2$, the rescaled zero-sum game condition is very stringent – it places exacting equality constraints on the elements of $A$ and $B$. However, for $n = 2$, every bimatrix game satisfies $a_{ij} + \delta_j = -cb_{ji} + \gamma_i$ for some $c$ potentially positive (in which case we have a rescaled zero-sum game) or negative (in which case the game is called a *partnership game*, and the coexistence equilibrium is unstable) (Hofbauer & Sigmund 1998). Thus, neutral oscillations arise whenever $c > 0$.

To see that this is true, we first suppose that $A$ and $B$ have the form

$$A = \begin{pmatrix} 0 & a_1 \\ a_2 & 0 \end{pmatrix} \qquad B = \begin{pmatrix} 0 & b_1 \\ b_2 & 0 \end{pmatrix} . \tag{S18}$$

If this is not the case, we can use constant column shifts to arrive at this form (e.g., in general, $a_1 = a_{12} - a_{22}$). Now consider the constants $c = -\frac{a_1+a_2}{b_1+b_2}$ and $\gamma_1 = \delta_1 = a_1 + cb_2$ and $\gamma_2 = \delta_2 = 0$. Examining the equation $a_{ij} + \delta_j - \gamma_i = -cb_{ji}$ for each $i$ and $j$, one verifies

$$\begin{aligned} 0 + \gamma_1 - \delta_1 &= 0 \\ a_1 + \gamma_2 - \delta_1 &= -cb_2 \\ a_2 + \gamma_1 - \delta_2 = -c(b_1 + b_2 - b2) &= -cb_1 \\ 0 &= 0 \end{aligned} \tag{S19}$$

and so the parameters $A$ and $B$ always constitute a rescaled zero-sum or partnership game. In the particular case of our model, $a_1+a_2 = -\alpha_{11}+\alpha_{21}+\alpha_{12}-\alpha_{22} = -I_s$ and $b_1+b_2 = -\beta_1-\beta_2$. $c$ is positive (as needed for cycles) when these signs disagree; since $b_1+b_2 = -\beta_1-\beta_2$ is always negative, $a_1 + a_2$ must be positive, meaning $I_s < 0$, as found by Bever *et al.* (1997).

### 7 Constants of motion

When $A$ and $B$ satisfy the rescaled zero-sum game condition, the function

$$H(\boldsymbol{p}, \boldsymbol{q}) = \sum_i p_i^\star \log p_i + c \sum_j q_j^\star \log q_j \tag{S20}$$

is a constant of motion for the dynamics (Hofbauer & Sigmund 1998). As above, we suppose that $A = -cB^T$, and shift the columns of each matrix as needed if this is not the case. Then consider the time derivative

$$\begin{aligned}
\frac{dH}{dt} &= \sum_i p_i^\star \frac{1}{p_i} \frac{dp_i}{dt} + c \sum_j q_j^\star \frac{1}{q_i} \frac{dq_i}{dt} \\
&= \sum_i p_i^\star \left( \sum_j \alpha_{ij}\, q_j - \sum_{j,k} \alpha_{jk}\, p_j\, q_k \right) + c \sum_j q_j^\star \left( \beta_i\, p_i - \sum_j \beta_j\, p_j\, q_j \right) \\
&= \sum_{i,j} \alpha_{ij}\, p_i^\star\, q_j - \sum_{j,k} \alpha_{jk}\, p_j\, q_k + c \sum_i \beta_i\, q_i^\star\, p_i - c \sum_j \beta_j\, p_j\, q_j \\
&= \sum_{i,j} \alpha_{ij}\, (p_i^\star - p_i)\, q_j + c \sum_i \beta_i\, (q_i^\star - q_i)\, p_i
\end{aligned}$$

Now, because $A = -cB^T$, we have

$$\begin{aligned}
&= c \sum_i \beta_i\, (-(p_i^\star - p_i)\, q_i + (q_i^\star - q_i)\, p_i) \\
&= c \sum_i \beta_i\, (-p_i^\star\, q_i + q_i^\star\, p_i)
\end{aligned}$$

and because $q_i^\star = p_i^\star = \frac{Z}{\beta_i}$, with $Z$ the normalizing constant,

$$\begin{aligned}
&= c\, Z \sum_i (-q_i + p_i) \\
&= 0
\end{aligned} \tag{S21}$$

In the last line, we use the fact that both sets of frequencies always sum to one.

Each orbit remains in the level set defined by the initial conditions, $(\boldsymbol{p}_0, \boldsymbol{q}_0)$:

$$H(\boldsymbol{p}_0, \boldsymbol{q}_0) = \sum_i p_i^\star \log p_i + c \sum_j q_j^\star \log q_j \tag{S22}$$

For the two-species model studied by Bever *et al.* (1997), these level sets precisely define the trajectories in the $(p, q)$ phase plane.

### 8 Equilibrium feasibility

Throughout this study, we focus primarily on the stability properties of the generalized Bever model. However, as mentioned above, coexistence also requires the existence of a feasible equilibrium – that is, an equilibrium where all frequencies are nonnegative. In the context of this model, feasibility is determined solely by the matrix $A$. If all elements of $A^{-1}\mathbf{1}$ share the same sign, the coexistence equilibrium is feasible. For even moderately large $n$, feasibility of the coexistence equilibrium is very unlikely if the parameters $\alpha_{ij}$ are iid random variables. However, the probability of feasibility has little bearing on the prospects for coexistence in this model. Even assuming the existence of a feasible equilibrium, our results show that robust coexistence of more than two species is impossible. To confirm that this is the case, we repeat the simulations shown in Fig. 2 (Main Text), but now rejecting parameter combinations that do not yield a feasible coexistence equilibrium. The results are show in Fig. S1. Conditioning on feasibility increases the probability that randomly parameterized two-species communities oscillate neutrally from $\frac{1}{4}$ to $\frac{1}{2}$, but has little effect on the results observed for $n > 3$. In particular, coexistence of more than two species is never observed, regardless of feasibility.

It is interesting to note that the rescaled zero-sum game condition, which ensures neutral stability of a fixed point, also ensures feasibility. This is easy to verify using the transformation explained in the section *Rescaled zero-sum games are neutrally stable*, above. Using column shifts applied to $A$ and $B$, one obtains a new system where both matrices are diagonal with constant signs. In other words, one finds a system of form $A = -cB^T$ with the same dynamics (and so the same equilibria) as the original. Because $B$ is a diagonal matrix, and we assume $\beta_i > 0$ for all $i$, both $\boldsymbol{p}^\star$ and $\boldsymbol{q}^\star$ will be feasible. However, we note that this property does not alter any of the conclusions of the Main Text. While the rescaled zero-sum game condition guarantees a weak form of coexistence (i.e. the existince of neutral oscillations), this behavior is extremely fragile; small changes in the model parameters will cause all but two species to go extinct.

#### 9 Adding frequency dependence

To illustrate the robustness of our main findings, we consider an extension of the Bever model to include direct intraspecific plant competition. Building on Eq. S1, we add a negative frequency-dependent term for each plant species:

$$\frac{dx_i}{dt} = x_i \left( \sum_j \alpha_{ij} q_j - c_i p_i \right) \tag{S23}$$

Here, $c_i$ specifies the strength of intraspecific competition. Soil dynamics remain exactly as in Eq. S1.

This model is conceptually close to the combined plant competition-feedback model introduced by Bever (2003). Unlike Bever, we consider only intraspecific plant interactions for simplicity. Additionally, while Bever took plant-plant interactions to be density-dependent, as in the Lotka-Volterra competition model, we assume frequency-dependent effects. As explained in the Main Text, this choice is motivated by consistency with the frequency-dependent nature of PSFs in this model.

The frequency dynamics associated with this model are given by

$$\begin{cases} \frac{dp_i}{dt} & = p_i \left( \sum_j \alpha_{ij}\, q_j - c_i\, p_i - \sum_j p_j (\sum_k \alpha_{jk}\, q_k - c_j\, p_j) \right) \;, \quad i = 1, \dots n \\ \frac{dq_i}{dt} & = q_i \left( \beta_i\, p_i - \sum_j \beta_j\, p_j\, q_j \right) \;. \end{cases} \tag{S24}$$

To consider small deviations from the canonical Bever model, we focus on the case where the negative frequency-dependence is weak relative to PSFs (i.e. $c_i$ parameters are much smaller than $\alpha_{ij}$ parameters). At the opposite extreme ($c_i \gg \alpha_{ij}$), it is easy to see that all plant species will coexist, with no meaningful role for PSFs. We also assume that frequency-dependence is equal for all plant species (i.e. $c_i = c$), for simplicity.

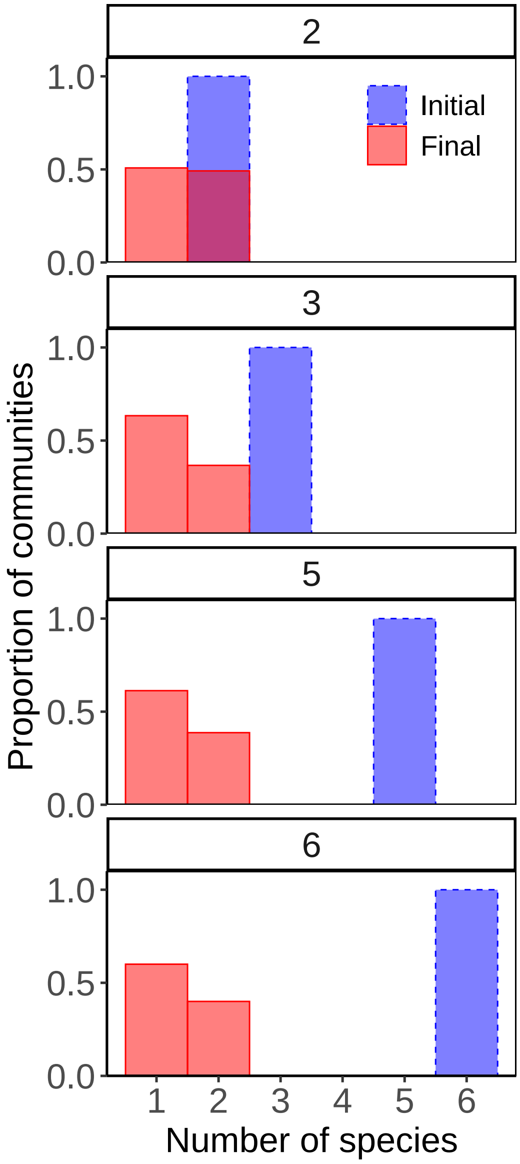


Figure 1: Final community sizes with varying initial richness, conditioned on feasible coexistence equilibrium. As in Fig. 2 (Main Text), except that parameter combinations yielding unfeasible equilibria were discarded. We continued sampling until 5000 feasible parameter sets were obtained for each level of initial richness. Conditioning on feasibility increases the probability that an initial community of two species coexists in a neutral cycle, but has negligible effect on the results for richer communities. In particular, coexistence of more than two species is never observed.

Now we study the stability properties of equilibria in this extended model. After some algebraic manipulations to remove the zero-sum constraints (as in the section *Local stability analysis*), we find that the community matrix for the coexistence equilibrium takes the form

$$J' = \begin{pmatrix} -cI & M_1 \\ M_2 & 0 \end{pmatrix} \tag{S25}$$

where

$$J = \begin{pmatrix} 0 & M_1 \\ M_2 & 0 \end{pmatrix} \tag{S26}$$

is the community matrix for the corresponding Bever model (i.e. the model with $c = 0$). We have already shown that the eigenvalues of $J$ must be of mixed signs or all purely imaginary. Let us denote those eigenvalues by $\lambda_i$. The eigenvalues of our extended matrix, which we call $\lambda'_i$, can be related to the $\lambda_i$ in a straightforward way. We first notice that the eigenvectors of $J'$ are closely related to the eigenvectors of $J$, which we write as $(\boldsymbol{u}_i, \boldsymbol{v}_i)^T$. The eigenvector equations for $J'$ take the form

$$\begin{pmatrix} -cI & M_1 \\ M_2 & 0 \end{pmatrix} \begin{pmatrix} \boldsymbol{u}_i \\ k_i \boldsymbol{v}_i \end{pmatrix} = \lambda'_i \begin{pmatrix} \boldsymbol{u}_i \\ k_i \boldsymbol{v}_i \end{pmatrix} \tag{S27}$$

with $k_i$ an undetermined constant. This system implies the relations $k_i \, \lambda'_i = \lambda_i$ and $\frac{\lambda'_i + c}{k_i} = \lambda_i$. Solving these equations for $\lambda'_i$ gives

$$\lambda'_i = \frac{-c \pm \sqrt{c^2 + 4\lambda_i^2}}{2} \tag{S28}$$

and finally, for small $c$, the approximation

$$\lambda'_i \approx \lambda_i - \frac{c}{2} \, . \tag{S29}$$

This analysis shows that there is a tight relationship between the stability properties of the Bever model and the extension with weak frequency-dependent self-regulation. If the underlying Bever model has an unstable coexistence equilibrium, where the eigenvalues $\lambda_i$ have mixed signs, then the extended model will have an unstable equilibrium as well. The

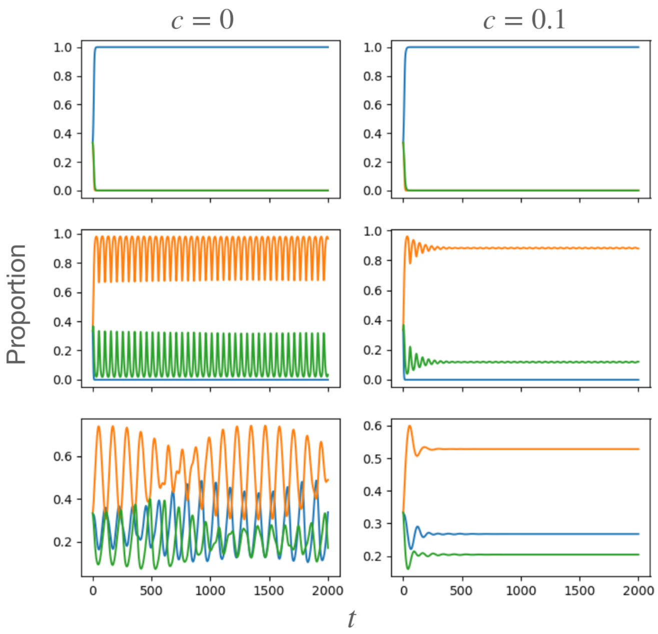


Figure 2: Representative dynamics for the extended Bever model with negative plant frequency-dependence. When the Bever model (left column) possess an unstable coexistence equilibrium, so will the extended model (right) with weak self-regulation (top row). On the other hand, when the Bever model possesses a neutrally stable equilibrium, the extended model will have a corresponding *stable* equilibrium, with the same number of species. We show an example where one of three species goes extinct and the other two cycle in Bever model, or stably coexist in the extended model (middle row). We also see that when the Bever model possess an $n$-species cycle (here $n = 3$), the extended model will have a stable equilibrium with all $n$ species. Such cases are only possible when the matrices $A$ and $B$ satisfy the rescaled zero-sum game condition, described in the Main Text and above.

slight shift by $\frac{c}{2}$ is not enough to push the positive real parts of these eigenvalues across zero, by assumption. The correspondence when all of the $\lambda_i$ are purely imaginary is more interesting. In this case, the eigenvalues of the extended model, $\lambda_i'$, will all have a small negative real part. This shift induces a qualitative change in the model dynamics: a neutrally stable equilibrium in the underlying Bever model becomes an asymptotically stable equilibrium in the model with frequency-dependence. Each of these cases is illustrated in Fig. S2.

Very weak frequency-dependence can only produce such a qualitative change when the underlying model is structurally unstable – i.e. when the real parts of the $\lambda_i$ are exactly zero. We have shown that this is only the case when the Bever model parameters meet the rescaled zero-sum game condition. Thus, even though the extended model can support stable coexistence, this outcome is subject to the same stringent conditions as are $n$-species oscillations in the Bever model. In particular, these parameterizations are never realized at random, and are not robust to small perturbations of the parameters.

This simple example demonstrates that the lack of robust $n$-species coexistence in the Bever model can be disentangled from the biologically unrealistic prediction of neutral oscillations. The generic behavior of the Bever model with more than two plant species is instability, and other ecological processes must be sufficiently strong to overcome this instability; very small modifications of the dynamics will not do.

#### 10 Numerical Simulations

To complement our analytical findings, we investigated the dynamics of many randomly parameterized communities using numerical simulations. In particular, we integrated Eq. S4 with 2, 3, 5, or 6 initial plant species and corresponding soil components. For each case, we sampled 5000 parameter sets at random and integrated the dynamics in Python using SciPy's (version 1.7.1) solve_ivp function with the "BDF" method. We sampled non-singular payoff matrices $A$ and $B$ with each non-zero element drawn independently from the uniform distribution U(0, 1). For every choice of parameters, we integrated the system until a subset with $\leq 2$ species was reached (which occurred in all cases). Code for reproducing all numerical simulations is available at `https://github.com/pablolich/plant_soil_feedback`.
